# Litter Microbiome Composition in Commercial Broiler Houses Under Contrasting Management: A Pilot Study

**DOI:** 10.1101/2025.10.31.685678

**Authors:** Elena G. Olson, Gaurav Gajurel, Steven C. Ricke, Brett Hale

## Abstract

Litter management affects broiler health, performance, and environmental impact, yet the effects of litter-applied amendments on the resident microbiome remain poorly characterized, particularly under commercial conditions. This pilot study evaluated the litter microbiome of three commercial broiler houses under contrasting management regimens. Two houses received a postbiotic litter amendment combined with sodium bisulfate (NaHSO₄) at 40% of the standard rate for one or two consecutive flocks, while a control house received the standard NaHSO₄ rate alone. Composite litter samples were collected following flock removal, and the V4 region of the 16S rRNA gene was sequenced to characterize prokaryotic community structure, diversity, and composition. Within-sample (α) diversity did not differ among houses (ANOVA, *p* > 0.05), whereas β diversity differed across Bray–Curtis, Jaccard, and weighted and unweighted UniFrac distances (PERMANOVA, *p* < 0.05). Pairwise comparisons detected no differences between the two amended houses, whereas each differed from the control across all distance metrics except weighted UniFrac for the house amended during a single flock (PERMANOVA, *q* < 0.05 for significant comparisons). Core microbiome and differential abundance analyses identified consistent compositional shifts, including greater representation of *Bifidobacterium* and lower relative abundance of *Staphylococcus* in both amended houses. Each management regimen was represented by a single house, so these observations should be interpreted as house-level associations rather than definitive treatment effects. Nevertheless, the consistency of microbiome responses between amended houses suggests that the defined alternative management regimen may influence resident microbial community composition in a manner relevant to litter conditioning, warranting further evaluation in replicated factorial designs.

## Introduction

Litter management is critical to broiler production, directly impacting bird health and welfare, environmental sustainability, and food safety. Poultry litter serves as a biologically complex substrate composed of fecal matter, feed residues, bedding material, and diverse microbial populations that interact with environmental and management factors. These resident microbial communities are shaped by variables such as stocking density, litter age and reuse, and the application of chemical or biological amendments (Bolan et al., 2010). As the poultry industry shifts toward antimicrobial stewardship and microbiome-informed production strategies, characterizing litter microbial ecology provides a foundation for evaluating and improving litter management practices (Olson et al., 2026; Ricke et al., 2026a).

Broiler producers apply acidifying amendments such as sodium bisulfate (NaHSO₄) to transiently reduce litter pH, limiting the conversion of nonvolatile ammonium (NH₄⁺) to atmospheric ammonia (NH₃) (Ritz et al., 2004; Rothrock et al., 2010). While the effectiveness of NaHSO₄ in mitigating NH₃ emissions is well established (Johnson et al., 2011), its effects on litter microbiome structure and composition remain less understood (Toppel et al., 2019; de Toledo et al., 2020; Johnson et al., 2021). For example, acidification can reduce total bacterial load (Johnson et al., 2021) but may favor the persistence of acid-tolerant bacteria, including *Salmonella* (Williams et al., 2012) and *Escherichia* (Joerger et al., 2020; Johnson et al., 2021). The potential for such community shifts following acidification highlights the need to investigate litter microbiome dynamics more holistically (e.g., via metabarcoding) and suggests that complementary interventions may be warranted to preserve microbiome stability while ensuring NH₃ reduction.

Biological interventions (e.g., pre-, pro-, post-, and synbiotics) are increasingly employed in poultry production systems to support bird health and production efficiency. These technologies are formulated primarily as feed or water additives (Callaway and Ricke, 2023) and have been shown to elevate mutualistic organisms in the alimentary tract, prime the host immune system, and improve production outcomes by enhancing nutrient uptake, digestibility, and intestinal integrity (Olson et al., 2025). However, the application of such interventions to litter remains comparatively underexplored relative to their use as ingestible interventions (Pedroso et al., 2013), and much of the existing literature reflects controlled microcosm conditions that may not capture microbial dynamics in commercial production environments (Johnson et al., 2021). Expanding biological intervention strategies to the litter environment is particularly relevant given that poultry litter harbors microbiota that directly impact bird health and litter physicochemistry, disperse into adjacent matrices (e.g., air, dust, and water lines), and contribute to the establishment of the chick alimentary tract microbiome (Gilchrist et al., 2007; Olson and Hale, 2026; Ricke et al., 2026b). Thus, biological amendments designed for litter application may offer an additional route to enhance food safety and bird performance.

The objective of this pilot study was to characterize the litter microbiome of commercial broiler houses under contrasting management regimens. Litter was sampled from a control house receiving an industry-standard NaHSO₄ rate and from two houses receiving a postbiotic litter amendment with reduced NaHSO₄ for one or two consecutive flocks. Prokaryotic communities were profiled by 16S rRNA gene sequencing and evaluated for within- and between-sample diversity, core microbiome membership, and differential abundance. These findings provide insight into how alternative litter management may influence the resident microbiome under commercial conditions and inform the design of future replicated studies.

## Materials and Methods

### Study design

The test site was in Sevier County, Arkansas, USA, and encompassed three commercial broiler houses with 500 X 43 ft dimensions (21,500 ft²; ∼1,997 m²), selected for their consistent history of litter management and bird productivity. The bedding material, which consisted of locally sourced rice (*Oryza sativa*) hulls, had been used for 18 consecutive broiler flocks prior to study initiation, with a cake layer removed between each flock. The houses were managed under a mechanical ventilation system with fans operating on a 300-s cycle for a minimum of 60 s at the beginning of the flock and increasing to 240 s by market weight (52 d). Temperature was initiated at 89–90 °F (31.7– 32.2 °C) and was gradually reduced to 65 to 68 °F (18.3–20.0 °C) as the birds approached market size, while relative humidity was maintained at approximately 50–70%. Feed composition followed a standard four-phase broiler regimen: Starter (d 1 to17), Grower (d 17–37), WD1/Finisher (d 36–47), and WD2/Finisher (d 48 onward). Two houses (denoted hereafter as “1” and “2”) received a postbiotic litter amendment (AgriGro Inc, Doniphan, MO, USA) at a rate of 48 oz/1,000 ft² (∼1.42 L/93 m²) 72 hrs before chick placement. According to the manufacturer, the amendment is formulated to reduce litter moisture and support broader litter conditioning. Treatment 2 also received the postbiotic during the preceding flock (**Table 1**). Postbiotic applications were made with a spot sprayer between litter windrowing and leveling.

**Table 1.**
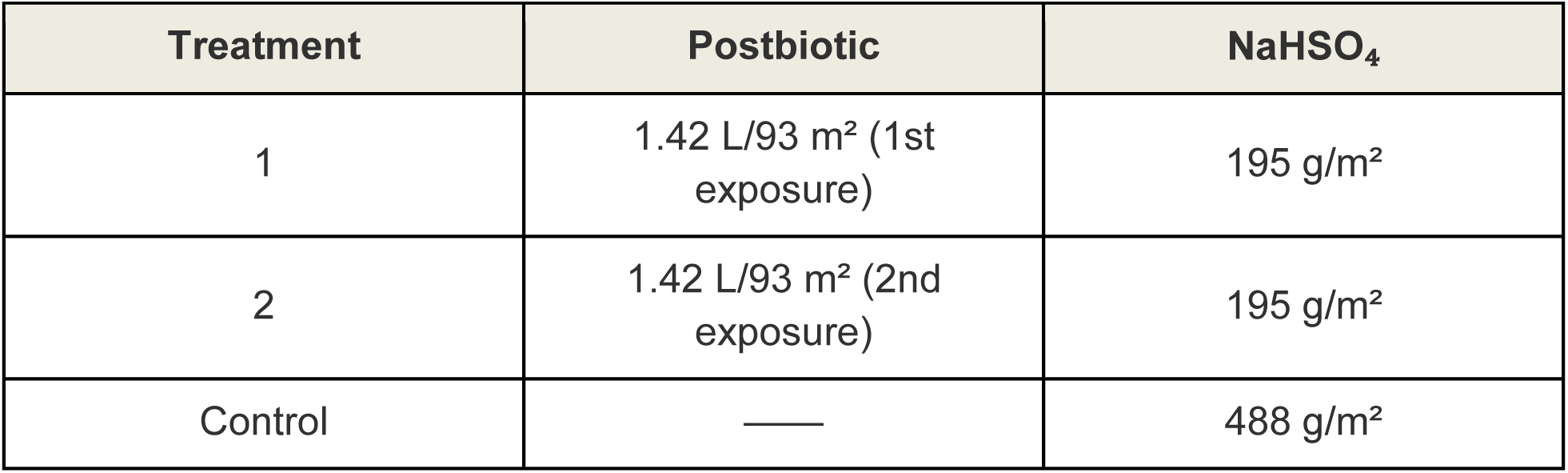
Litter amendment regimen for each treatment.

| Treatment | Postbiotic | NaHSO₄ |
|---|---|---|
| 1 | 1.42 L/93 m² (1st exposure) | 195 g/m² |
| 2 | 1.42 L/93 m² (2nd exposure) | 195 g/m² |
| Control | — | 488 g/m² |

Approximately 16,333 Ross 308 chicks (Aviagen, Huntsville, AL, USA) were placed per house, corresponding to a stocking density of 1.32 birds/ft² (8.18 birds/m²). Less than 24 hours prior to chick placement, an NaHSO₄-based litter amendment (Jones-Hamilton Co., Maumee, OH, USA) was applied to houses 1 and 2 at a rate of 40 lbs/1,000 ft² (∼195 g/m²) and to house 3 (“Control”) at 100 lbs/1,000 ft² (∼488 g/m²) (**Table 1**). All other production parameters remained constant across houses.

### Sample collection and DNA isolation

On the day following bird removal, six composite litter samples (*n* = 18) were collected from each house to capture spatial heterogeneity (Locatelli et al., 2017). Each composite consisted of five subsamples collected to a depth of 10 cm using a sanitized trowel, with a new trowel used for each house to prevent cross-contamination. Subsamples were pooled into sterile ziplock bags and immediately flash-frozen on dry ice. Samples were then shipped on dry ice to the University of Missouri (MU) Genomics Technology Core (Columbia, MO, USA), where they were stored at −80 °C until DNA isolation. Because each regimen was applied to a single house, the composite samples represent spatial subsamples within houses rather than independent replicates of each regimen.

DNA was isolated using the QIAamp PowerFecal Pro DNA Kit (Cat. #51804; Qiagen, Hilden, Germany) according to the manufacturer’s instructions, with samples homogenized using a TissueLyser II (Qiagen) at 30 Hz for 10 minutes in place of the vortex adapter specified in the protocol. DNA concentrations were determined fluorometrically using a Qubit 2.0 fluorometer (Thermo Fisher Scientific, Waltham, MA, USA) with the Quant-iT dsDNA Broad-Range Assay Kit (Cat. #Q33130) and subsequently normalized to a consistent concentration and volume.

### 16S rRNA gene library preparation and sequencing

Amplicons were generated by targeting the V4 region of the 16S rRNA gene using universal primers U515F/806R, flanked by Illumina standard adapter sequences (Caporaso et al., 2011; Walters and Caporaso et al., 2011). All reactions employed dual-indexed forward and reverse primers. PCR was performed in 50 µL reactions containing 100 ng DNA, 0.2 µM of each primer, 200 µM of each dNTP, 1 U of Phusion High-Fidelity DNA Polymerase (Cat #F530S; Thermo Fisher Scientific), and the appropriate volume of 5X Phusion HF Buffer (Cat. #F518L; Thermo Fisher Scientific). Amplification conditions included initial denaturation at 98 °C for 3 min, followed by 25 cycles of 98 °C for 15 s, 50 °C for 30 s, and 72 °C for 30 s, with a final extension at 72 °C for 7 min.

Following amplification, 5 µL of product from each reaction was pooled, mixed thoroughly, and purified using Axygen Axyprep MagPCR Clean-up Beads (Cat. #14-223-227; Thermo Fisher Scientific) at a 1:1 bead-to-amplicon volume ratio. The mixture was incubated for 15 min at room temperature, washed twice with 80% ethanol, and air-dried. The purified pellet was resuspended in 32.5 µL of EB buffer (Cat. #19086; Qiagen), incubated at room temperature for 2 min, and placed on a magnetic stand for 5 min to recover the supernatant.

The final pooled amplicon library was evaluated using the Advanced Analytical Fragment Analyzer (Agilent Technologies, Santa Clara, CA, USA), quantified with the Quant-iT High Sensitivity dsDNA Assay Kit (Cat. #Q33120; Thermo Fisher Scientific) and diluted according to Illumina’s protocol for sequencing on the MiSeq platform as 2 × 250 bp paired-end reads.

### Bioinformatic analysis

DNA sequences were assembled and annotated at the MU Bioinformatics and Analytics Core (Columbia, MO, USA). Adapter and primer removal was conducted using Cutadapt (v2.6) (Martin, 2011). The forward primer was trimmed from the 5′ end of the forward read, and, if present, the reverse complement of the reverse primer was removed from the 3′ end of the same read along with all downstream bases, resulting in bidirectional trimming if the insert was shorter than the amplicon. The reverse read was processed analogously, with primer roles reversed. Read pairs were discarded if either read failed to match a 5′ primer, with an allowed error rate of 0.1 and a minimum overlap of 3 bp at the 3′ end of the primer for removal. Each read underwent two passes to ensure complete primer removal.

Amplicon sequence variant (ASV) inference was performed using the DADA2 plugin (v1.30.0) (Callahan et al., 2016) within QIIME2 (v2024.5.1) (Bolyen et al., 2019). Denoising parameters included truncating both forward and reverse reads to 150 bases, discarding reads with more than 2.0 expected errors, and removing chimeras using the ‘consensus’ method. The QIIME2 environment incorporated Python (v3.9.19) and BIOM (v2.1.15) for downstream processing. Taxonomic classification was performed using the ‘classify-sklearn’ method with the SILVA v138.1 reference database (Quast et al., 2012).

### Statistical analysis

The QIIME2 artifacts were imported into R (v4.4.0) (R Core Team, 2024) within RStudio (Posit team, 2024) with the qiime2R package (v0.99.6) (Bisanz, 2018) and were used to create a “phyloseq” S4 object (v1.48.0) (McMurdie and Holmes, 2013). One Control house sample with <1,000 reads was removed prior to analysis. Sequencing depth was subsequently estimated using Good’s coverage, calculated with the ‘phyloseq_coverage’ function from the metagMisc package (v0.5.0) (Mikryukov, 2020) with the formula *1 - (F1/N)*, where *F1* is the number of singleton ASVs and *N* is the read sum for all ASVs (Good, 1953; Good and Toulmin, 1956; Chao and Jost, 2012). Observed ASVs (richness) were evaluated as a function of sequencing depth using the ‘rarecurve’ function from the vegan package (v2.6.1) (Dixon, 2003). All visualizations were generated with ggplot2 (v3.4.2) (Wickham and Sievert, 2009). The rooted phylogenetic tree was converted to a “phylo” object using the ape package (v5.8) (Paradis et al., 2004) and visualized as a cladogram with ggtree (v3.12.0) (Yu et al., 2017).

Within-sample (α) diversity was calculated after rarefying all samples to the minimum sequencing depth (170,641 reads per sample). Richness and Shannon diversity (Shannon, 1948) were determined with the ‘estimate_richness’ phyloseq function, Pielou’s evenness (Pielou, 1966) with the ‘evenness’ function from the microbiome package (v1.26.0) (Lahti and Shetty, 2018), and Faith’s phylogenetic diversity (Faith, 1992) with the ‘pd’ function in picante (v1.8.2) (Kembel et al., 2010). For each metric, Levene’s test (Levene, 1960) was used to assess homogeneity of variance across treatments via the ‘leveneTest’ function in the car package (v3.1-2) (Fox et al., 2012), and the Shapiro–Wilk test (Shapiro and Wilk, 1965) was used to evaluate normality within each treatment group with ‘shapiro.test’ from the stats package (v4.4.0). All assumptions of homogeneity and normality were met (*p* > 0.05), and differences among houses were evaluated using analysis of variance (ANOVA) models fitted with the ‘aov’ function in stats. The α diversity estimates were visualized with ggplot2.

Prior to β diversity analysis, reads were normalized using cumulative sum scaling (CSS) with the ‘cumNormStat’ and ‘cumNorm’ functions from the metagenomeSeq package (v1.46.0) (Paulson et al., 2013). Global dissimilarity matrices were then constructed using Bray–Curtis (Bray and Curtis, 1957), Jaccard (1908), Weighted UniFrac, and Unweighted UniFrac (Lozupone and Knight, 2005) distances via the ‘distance’ function in phyloseq. Ordination was performed using Principal Coordinates Analysis (PCoA) with the ‘ordinate’ function in phyloseq, and results were visualized with ggplot2. Compositional dissimilarity among treatment levels was assessed statistically using permutational multivariate analysis of variance (PERMANOVA) with the ‘adonis2’ function from vegan (v2.6-6.1) with 9,999 permutations. As PERMANOVA identified significant differences (*p* < 0.05) for all distance metrics, multilevel pairwise comparisons were performed using ‘pairwise.adonis’ from the pairwiseAdonis package (v0.4.1) (Martinez Arbizu, 2020), applying false discovery rate (FDR) correction to control for multiple testing. Homogeneity of multivariate dispersions was evaluated with the ‘betadisper’ function from vegan, and distances of samples to their respective group spatial medians were compared using conventional ANOVA via the ‘anova’ function. Pairwise PERMANOVA R^2^ values were visualized with the ComplexHeatmap package (v2.20.0) (Gu et al., 2016).

Global and treatment-specific core microbiome features were defined as ASVs present in all but one sample per condition, following the “relaxed core” concept proposed by Mejia et al. (2025). Per-sample relative abundances of core ASVs were calculated from CSS-normalized counts using the ‘transform_sample_counts’ function in phyloseq and visualized with the ComplexHeatmap package. The top 10 most represented genera, families, and orders by ASV count within each core microbiome were identified and visualized using ggplot2. Shared and exclusive ASVs among core groups were further visualized using the ggvenn package (v0.1.10) (Yan, 2021). Core microbiomes were clustered using Jaccard distance, calculated with the ‘vegdist’ function from the vegan package and visualized in ggplot2.

Features differentially abundant among treatment levels were identified using the MaAsLin2 package (Microbiome Multivariable Associations with Linear Models) (v1.18.0) (Mallick et al., 2021). Log-transformed linear models were constructed using total sum scaling (TSS)-normalized count data following MaAsLin2 default parameters. Pairwise treatment comparisons were performed independently. To reduce the influence of spurious or environmentally stochastic features in the absence of a shared baseline, analyses were restricted to features present in the core microbiomes of at least two treatment groups. Genus-, family-, and order-level counts were additionally generated using the ‘tax_glom’ function from the phyloseq package and reanalyzed using the same MaAsLin2 parameters. Features with a Benjamini–Hochberg-adjusted *p*-value (*q* < 0.25) (Benjamini and Hochberg, 1995) were considered differentially abundant as recommended by MaAsLin2 documentation (Mallick et al., 2021). The ASV-level results were visualized using volcano plots of MaAsLin2 coefficients against −log₁₀(*q*-values) in *ggplot2*. At higher levels, MaAsLin2 coefficients were visualized using the ComplexHeatmap package. Shared and unique features across pairwise comparisons were visualized with the ggvenn package.

## Results

### Sequencing overview and taxonomic composition of the litter microbiome

The objective of the present study was to characterize the litter microbiome of commercial broiler houses under contrasting litter management regimens. Comparisons were made between a house receiving the postbiotic amendment combined with NaHSO₄ at 40% of the recommended rate for the first time (treatment 1), a house treated with the same combination for a second consecutive flock (treatment 2), and a control house receiving only the recommended NaHSO₄ application. Six composite litter samples were collected per house, and the V4 region of the 16S rRNA gene was amplified and sequenced to assess prokaryotic microbiome differences among houses.

Illumina V4 sequencing yielded 333,596 ± 48,470 reads per sample (mean ± 1 SD), with Good’s coverage estimates (0.999994 ± 6e–6) and rarefaction curves supporting adequate sampling depth (**Figure 1a–c**). Denoising yielded 2,938 ASVs, with individual samples containing 531.94 ± 157.88. The dominant phyla were Firmicutes (59.87% of reads, 69.8% of ASVs), Actinobacteriota (35.95% of reads, 11.37% of ASVs), and Proteobacteria (3.31% of reads, 7.26% of ASVs). The most abundant classes were Bacilli (54.58% of reads, 25.61% of ASVs), Actinobacteria (35.94% of reads, 10.51% of ASVs), and Clostridia (5.27% of reads, 43.48% of ASVs). Dominant orders included Staphylococcales (30.48% of reads, 4.16% of ASVs), Micrococcales (19.80% of reads, 5.92% of ASVs), and Lactobacillales (17.74% of reads, 7.84% of ASVs). At family level, Staphylococcaceae (30.48% of reads, 4.16% of ASVs), Dermabacteraceae (10.01% of reads, 1.10% of ASVs), and Lactobacillaceae (9.40% of reads, 3.30% of ASVs) were most abundant (**Figure 1d**). The rooted phylogenetic tree showed that ASVs assigned to the dominant bacterial families generally clustered within related lineages (**Figure 1e**).

**Figure 1.**
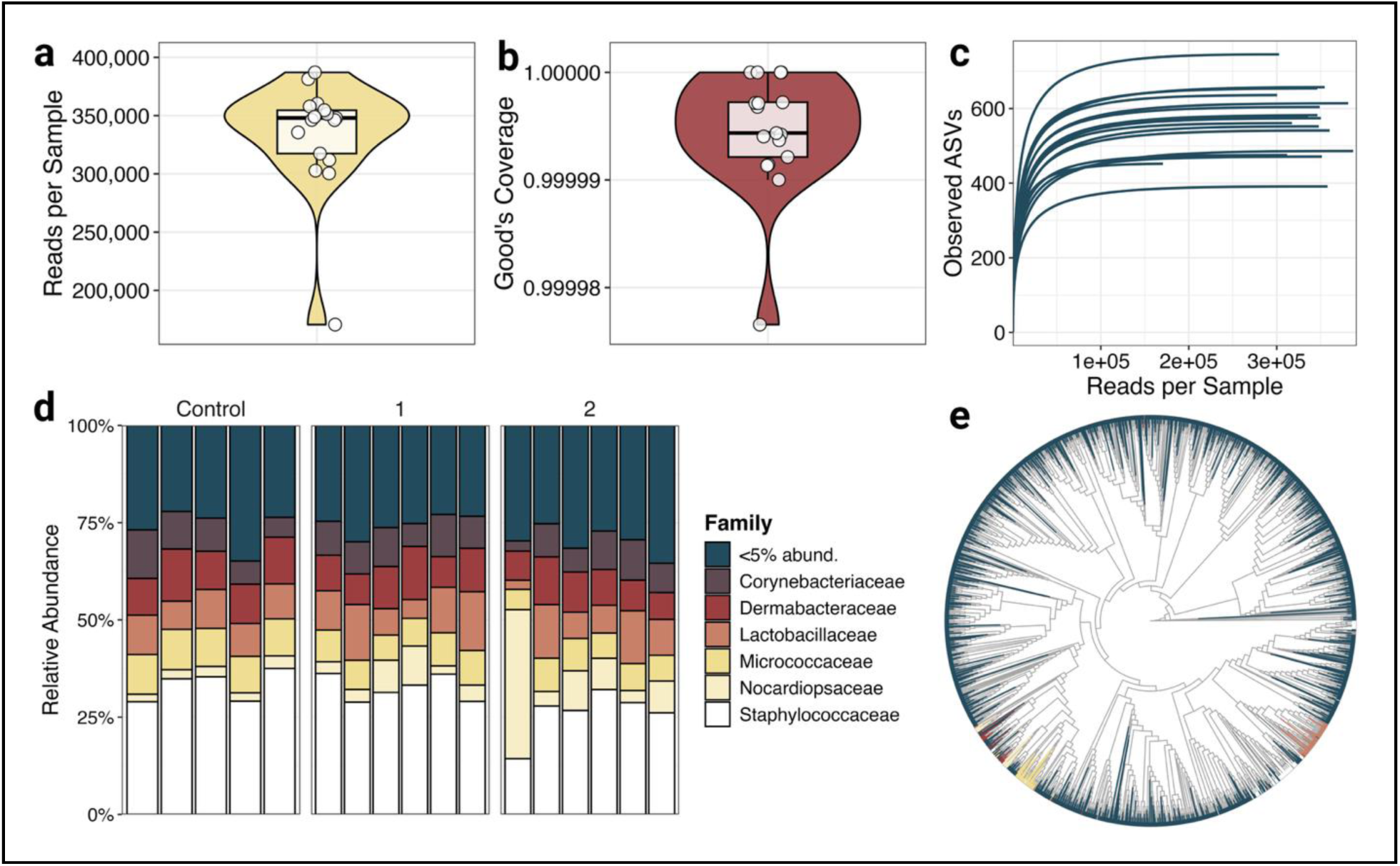
Overview of the poultry litter microbiome. (**a**) Sequencing depth distribution. (**b**) Distribution of Good’s coverage determined with the formula: *1 - (F1/N)*, where *F1* is the number of singleton ASVs and N is the read sum for all ASVs. (**c**) Rarefaction curve showing observed ASVs (richness) as a function of sequencing depth (reads per sample). (**d**) Stacked bar plot of family-level per-sample composition. Families contributing less than 5% of total read abundance across all samples were grouped into “<5% abund.” after agglomerating counts to the Family level. (**e**) Rooted phylogenetic tree. Tip color corresponds to family utilizing the same key as the stacked bar plot.

### Microbiome composition differed between amended and control houses

Alpha (α) diversity was evaluated for Faith’s phylogenetic diversity, observed ASVs, Shannon diversity, and Pielou’s evenness using libraries rarefied to the minimum sequencing depth (170,641 reads per sample). No significant differences were detected among houses for any metric (ANOVA, *p* ≥ 0.68) (**Figure 2a**). Count matrices were then normalized using CSS, and β diversity was evaluated using Bray–Curtis, Jaccard, weighted UniFrac, and unweighted UniFrac distances. Principal coordinate analyses illustrated differences in community composition among houses (**Figure 2b**), with significant overall differences detected across all four distance metrics (PERMANOVA, *p* < 0.05).

**Figure 2.**
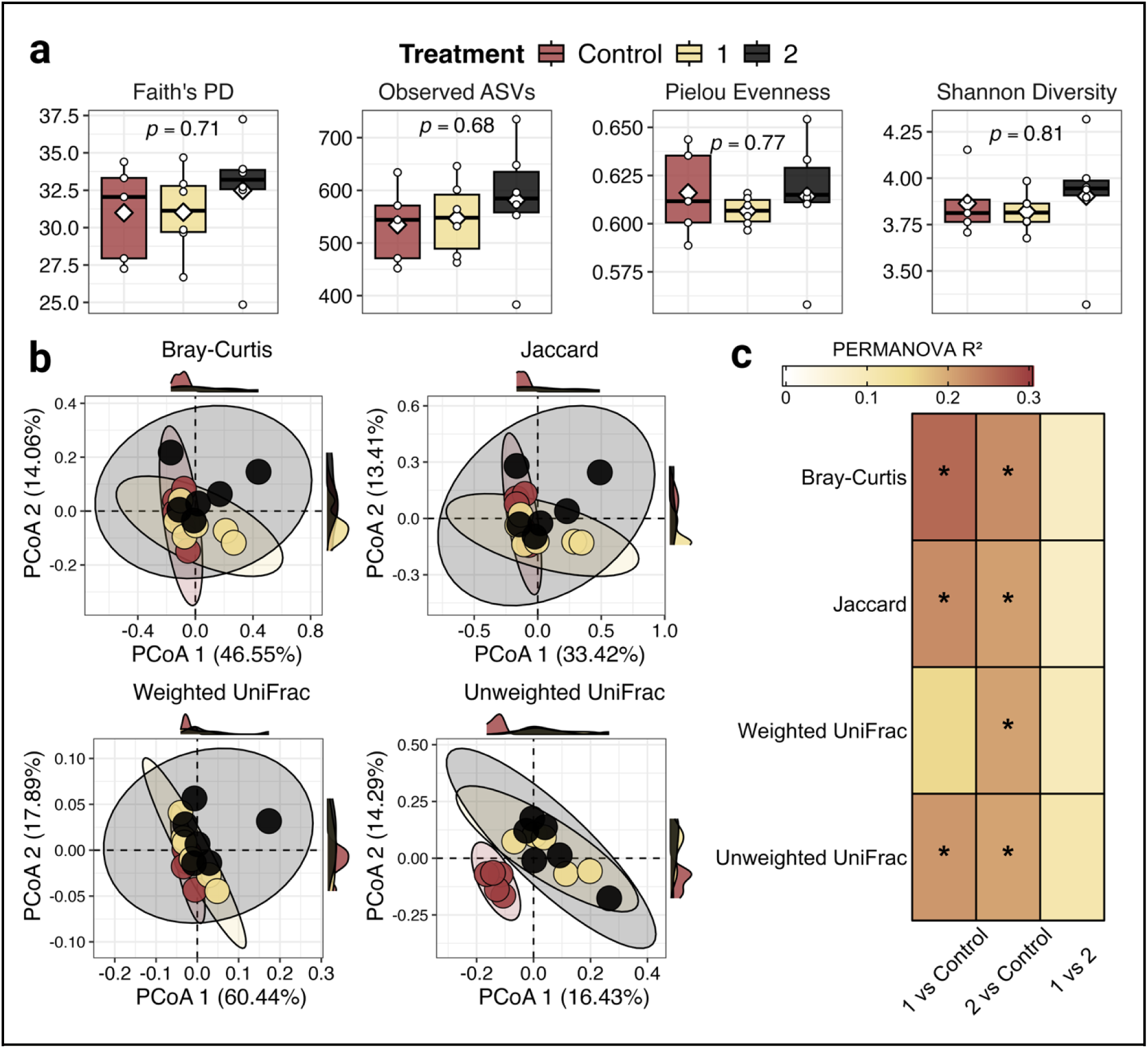
α and β diversity of the poultry litter microbiome across contrasting litter management regimens. (**a**) Boxplots showing Faith’s phylogenetic diversity, observed ASVs, Pielou’s evenness, and Shannon diversity. Individual points represent samples, white diamonds indicate means, and *p*-values correspond to ANOVA comparisons among houses. (**b**) PCoA ordinations based on Bray–Curtis, Jaccard, weighted UniFrac, and unweighted UniFrac distances. Shaded ellipses represent group distributions, and marginal density plots show sample distributions along each axis. (**c**) Heatmap of pairwise PERMANOVA results across the four distance metrics. Cell colors represent R² values, and asterisks indicate significant differences following FDR correction (*q* < 0.05).

Pairwise comparisons detected differences between treatment 1 and the control across all distance metrics except weighted UniFrac (*q* = 0.29, R² = 0.15), whereas treatment 2 differed from the control across all four metrics (*q* < 0.05) (**Figure 2c**). No significant differences were detected between treatments 1 and 2 (*q* ≥ 0.14). Global tests of β dispersion were nonsignificant across all distance metrics (ANOVA, *p* ≥ 0.30), providing no evidence that differences in within-house dispersion accounted for the observed compositional differences.

### Core microbiome membership differed among houses

A total of 170 ASVs were identified in the global core microbiome (**Figure 3a, d**). The most represented genus was *Christensenellaceae_R-7_group* (6 ASVs), followed by eight genera represented by four ASVs each (**Figure 3c**). At the family level, Lachnospiraceae (19 ASVs), Bacillaceae (16 ASVs), and Oscillospiraceae (14 ASVs) contained the greatest numbers of core ASVs, while the most represented orders were Oscillospirales (31 ASVs), Lactobacillales (25 ASVs), and Micrococcales (21 ASVs) (**Figure 3c**).

**Figure 3.**
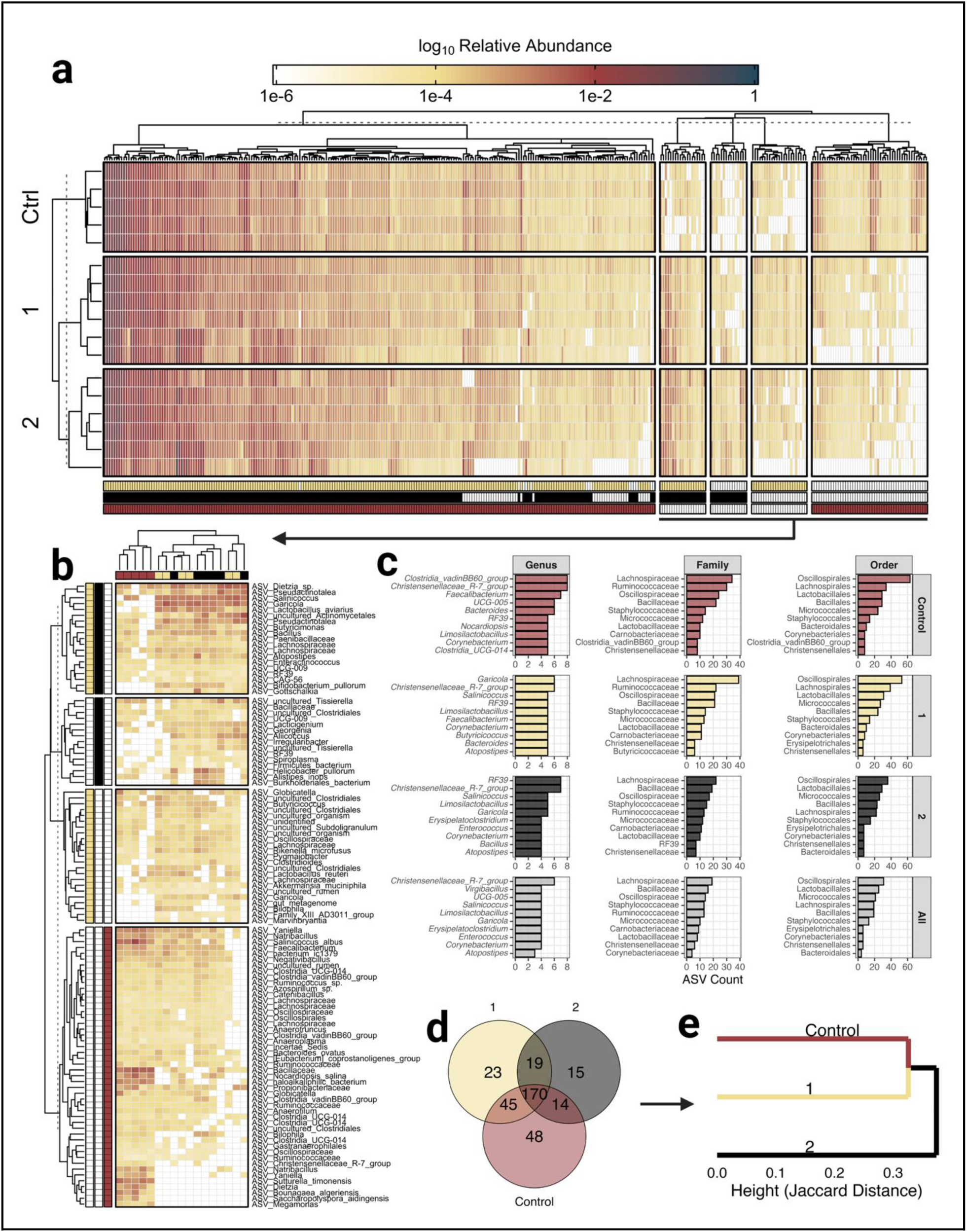
Global and treatment-specific core microbiome analysis. (**a**) Relative abundance (log₁₀) heatmap of ASVs represented in one or more core microbiomes. Column annotations reflect representation in control (red), 1 (yellow), and 2 (black) treatments. (**b**) Relative abundance heatmap of ASVs represented exclusively in control, 1, 2, or 1 and 2 (yellow and black row annotations). (**c**) Top ten genera, families, and orders represented in global and treatment-specific core microbiomes. (**d**) Venn diagram depicting shared and exclusive ASV counts across the treatment-specific core microbiomes. (**e**) Clustering of treatment-specific core microbiomes with Jaccard distances.

Characterization of the house-specific core microbiomes identified 334 distinct ASVs, comprising the 170 shared globally and 164 present in only one or two house cores (**Figure 3a–b, d**). Treatment 1 shared 215 core ASVs with the control, while treatment 2 shared 184 with the control. The two amended houses shared 189 core ASVs (**Figure 3d**). Forty-eight ASVs were unique to the control core microbiome, including ASVs classified as *Faecalibacterium* and *Nocardiopsis salina*, compared with 23 and 15 unique ASVs in treatments 1 and 2, respectively (**Figure 3b**). Nineteen ASVs were shared between the two amended houses but absent from the control, including ASVs classified as *Lactobacillus aviarius* and *Bifidobacterium pullorum* (**Figure 3b**). Jaccard-based clustering of core ASV membership indicated greater dissimilarity between the control and treatment 2 than between the control and treatment 1 (**Figure 3e**).

Genus-level representation within the core microbiomes also varied among houses. In the control, *Clostridia_vadinBB60_group* and *Christensenellaceae_R-7_group* were most represented (8 ASVs each), followed by *Faecalibacterium* (7 ASVs), *UCG-005* (6 ASVs), and *Bacteroides* (6 ASVs) (**Figure 3c**). Treatment 1 was characterized by *Garciella* and *Christensenellaceae_R-7_group* (6 ASVs each), while treatment 2 had the greatest representation of *Christensenellaceae_R-7_group* and *RF39* (7 ASVs each). Despite these genus-level differences, the house-specific core microbiomes exhibited broadly similar taxonomic representation at the family and order levels (**Figure 3c**).

### Differential abundance analysis identified consistent differences between amended and control houses

Differential abundance analysis was performed at the ASV level using log- transformed linear models, with the control house specified as the reference. A total of 115 ASVs were differentially abundant (*q* < 0.25) in one or both amended houses relative to the control. Of these, 82 were identified in treatment 1 (39 enriched, 43 depleted) and 81 in treatment 2 (45 enriched, 36 depleted) (**Figure 4a**). Forty-eight ASVs (41.74%) were shared between comparisons, all exhibiting consistent directional changes (23 enriched, 25 depleted) (**Figure 4a**).

**Figure 4.**
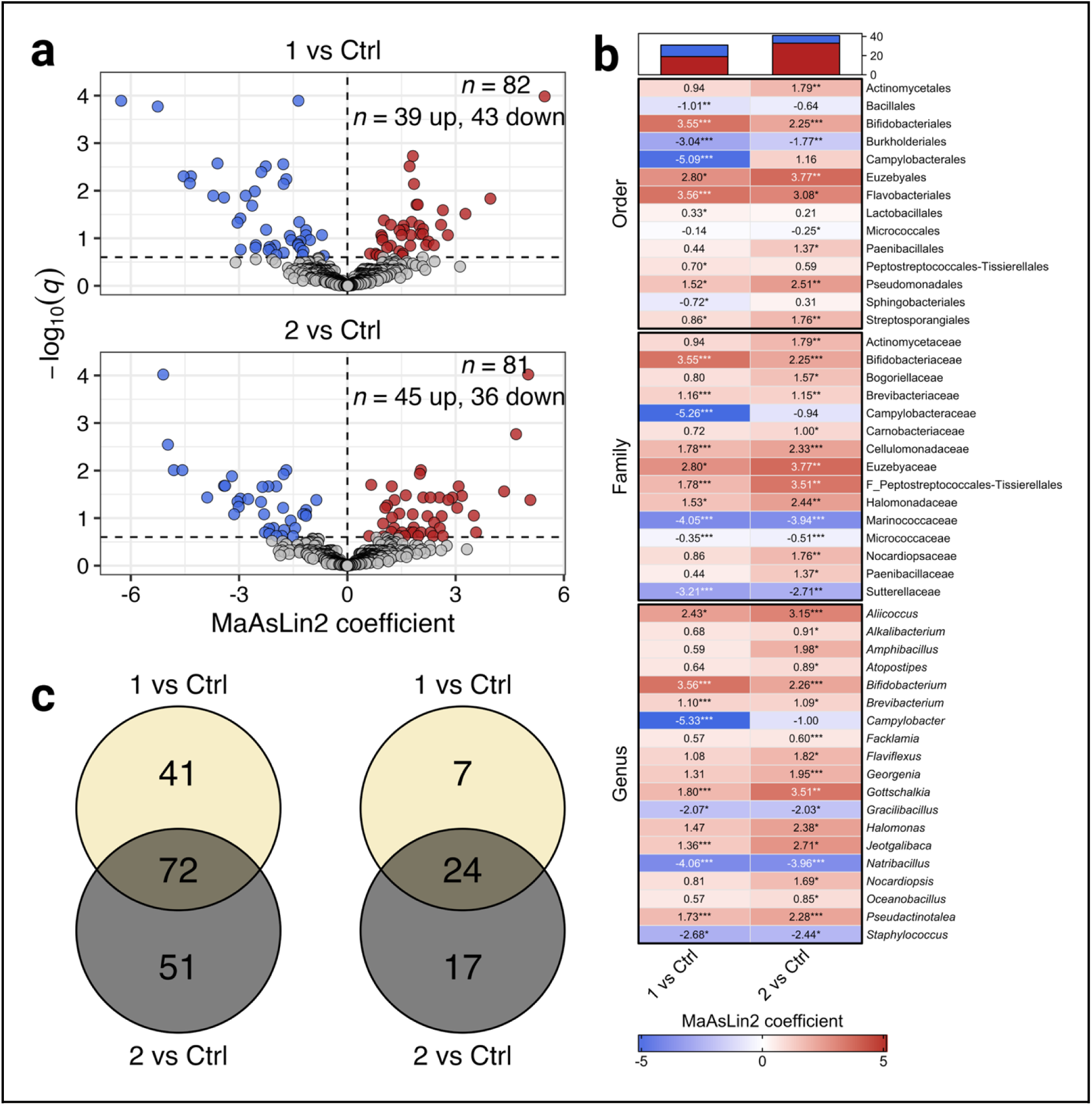
Differential abundance of the poultry litter microbiome across contrasting litter management regimens. (**a**) ASV-level volcano plots comparing each amended house with the control. Points represent individual ASVs, with red and blue indicating enriched and depleted features, respectively. Dashed horizontal lines indicate the exploratory FDR threshold (*q* < 0.25). (**b**) Heatmap of MaAsLin2 coefficients at the order, family, and genus levels. Cell colors indicate the direction and magnitude of associations, with red representing enrichment and blue representing depletion relative to the control. Asterisks indicate FDR-adjusted significance (*** *q* < 0.05, ** *q* < 0.10, * *q* < 0.25). The stacked bar plots summarize enriched and depleted features meeting *q* < 0.25. (**c**) Venn diagrams showing overlap in differentially abundant features between amended-versus-control comparisons, including all taxonomic levels (left; ASV through order) and genus-, family-, and order-level features (right).

Counts were subsequently agglomerated at the genus, family, and order levels and reanalyzed using the same MaAsLin2 parameters. At the genus level, 20 features were differentially abundant in one or both amended houses relative to the control (**Figure 4b**). Ten were identified in treatment 1 (6 enriched, 4 depleted) and 19 in treatment 2 (16 enriched, 3 depleted), with nine shared between comparisons (6 enriched, 3 depleted). Genera enriched in both amended houses included *Bifidobacterium* (treatment 1: coef = 3.56, *q* = 0.04; treatment 2: coef = 2.26, *q* = 0.01) and *Brevibacterium* (treatment 1: coef = 1.10, *q* = 0.04; treatment 2: coef = 1.09, *q* = 0.14), whereas *Staphylococcus* was consistently depleted (treatment 1: coef = −2.68, *q* = 0.16; treatment 2: coef = −2.44, *q* = 0.11). Differential abundance of *Campylobacter* was detected only in treatment 1 relative to the control (coef = −5.33, *q* = 9 × 10⁻⁵), while treatment 2 exhibited additional enrichment of *Atopostipes* (coef = 0.89, *q* = 0.23), *Halomonas* (coef = 2.38, *q* = 0.11), *Nocardiopsis* (coef = 1.69, *q* = 0.14), *Oceanobacillus* (coef = 0.85, *q* = 0.14), and nine other genera (**Figure 4b**).

Family-level analysis identified 15 differentially abundant features, including 10 in treatment 1 (6 enriched, 4 depleted) and 14 in treatment 2 (11 enriched, 3 depleted). Nine families (60%) were shared between comparisons, with consistent directional changes (6 enriched, 3 depleted), largely reflecting genus-level patterns (**Figure 4b**). These included enrichment of Bifidobacteriaceae, Brevibacteriaceae, and four additional families, alongside depletion of Marinococcaceae, Micrococcaceae, and Sutterellaceae in both amended houses. Campylobacteraceae was depleted only in treatment 1 (coef = −5.26, *q* = 5.3 × 10⁻⁵), whereas Actinomycetaceae, Bogoriellaceae, Carnobacteriaceae, Nocardiopsaceae, and Paenibacillaceae were enriched only in treatment 2 (**Figure 4b**). At the order level, 14 features were differentially abundant across comparisons, with 11 in treatment 1 (7 enriched, 4 depleted) and nine in treatment 2 (7 enriched, 2 depleted). Six orders (42.86%) were shared, showing consistent directional changes (5 enriched, 1 depleted) and generally reflecting patterns observed at lower taxonomic ranks (**Figure 4b**).

Overall, the two amended houses exhibited partially concordant differential abundance profiles relative to the control. Twenty-four of 48 (50%) genus-to-order features and 72 of 164 (43.9%) total features were shared between comparisons, with all 72 exhibiting consistent directional changes (**Figure 4c**). The remaining features were identified exclusively in treatment 1 (41/164, 25%) or treatment 2 (51/164, 31.1%) relative to the control, indicating additional house-specific differences in microbiome composition (**Figure 4c**).

## Discussion

Litter amendments are typically evaluated by their effects on atmospheric NH₃ emission, moisture, and bird performance (de Toledo et al., 2020), whereas their influence on the litter microbiome has received less attention. In the present study, litter prokaryotic microbiome composition in houses receiving a postbiotic with reduced NaHSO₄ differed from that of a house receiving a standard NaHSO₄ rate, without differences in α diversity. Do et al. (2024) similarly reported minimal effects of an organic litter amendment on α diversity, whereas Johnson et al. (2021) observed increased richness shortly after NaHSO₄ application but little change in community structure unless the litter was also top-dressed with fresh bedding. Together, these findings suggest that litter amendments may act more on community membership and relative abundance than on overall diversity. The two amended houses also did not differ significantly across any distance metric, indicating similar community composition following one and two consecutive applications.

The core microbiome is thought to encompass functionally important community members that interact synergistically or antagonistically with transient taxa (Neu et al., 2021; Jiao et al., 2022). In the present study, *Christensenellaceae_R-7_group* was the most represented genus-level group in the global core. This taxon is frequently detected in the gastrointestinal tract of livestock, including broilers (Shi et al., 2023), consistent with fecal contributions to the litter microbiome and the bidirectional exchange of microorganisms between litter and the alimentary tract (Cressman et al., 2010; Olson and Hale, 2026). Although its abundance has been associated with improved feed efficiency in broilers (He et al., 2023), negative associations with body weight and average daily gain have also been reported (Xu et al., 2025), suggesting that its ecological significance may be context-dependent.

While substantial core membership was shared among houses, distinct differences were observed in house-specific core ASVs, particularly between the control and treatment 2. Several ASVs exclusive to the amended-house cores belonged to groups containing members previously associated with probiotic applications or improved poultry performance, including *Bifidobacterium*, Paenibacillaceae, *Bacillus*, and *Butyricimonas* (Grant et al., 2018; Wu et al., 2019; Halder et al., 2024; Kairmi et al., 2024). *Salinicoccus* and *Atopostipes*, both reported in poultry litter compost, were also more represented in the amended-house cores (Kubasova et al., 2022; Wang et al., 2022). Conversely, the control core uniquely contained three Clostridia UCG-014 ASVs and greater representation of *Faecalibacterium*. Clostridia UCG-014 has been reported to be significantly enriched following *Campylobacter jejuni* challenge in broilers (Valečková et al., 2023), whereas *Faecalibacterium* has been reported to increase following NaHSO₄ application in poultry litter (Johnson et al., 2021). These differences suggest potentially relevant ecological variation in core membership, although the functional significance of individual taxa and their associations with litter management remain to be established.

Differential abundance analysis complemented the core microbiome findings by identifying taxonomic associations that were consistent between the amended houses relative to the control. Both amended houses exhibited higher relative abundance of *Bifidobacterium*, *Brevibacterium*, *Gottschalkia*, *Jeotgalibaca*, and *Pseudactinotalea*. Among these, *Brevibacterium* is a common member of the poultry litter microbiome associated with organic matter decomposition and litter maturation (Dumas et al., 2011; Kubasova et al., 2022), while *Gottschalkia* is a purinolytic genus capable of degrading uric acid, the predominant nitrogenous compound in poultry excreta (Rothrock et al., 2010; Poehlein et al., 2017). Given that NaHSO₄ acidification can suppress microbial nitrogen mineralization, the greater representation of *Gottschalkia* in both amended houses may reflect conditions more favorable to uric acid degradation under reduced acidifier application. *Bifidobacterium*, which was also represented in the amended-house cores, includes members associated with poultry intestinal health and widely used in probiotic formulations (Halder et al., 2024).

Additional taxonomic differences were identified in the house receiving two consecutive postbiotic applications, including higher relative abundance of *Alkalibacterium*, *Oceanobacillus*, *Facklamia*, *Flaviflexus*, *Halomonas*, *Atopostipes*, and *Nocardiopsis*. Several of these genera have previously been reported in reused poultry litter or litter compost (Lu et al., 2003; Johnson et al., 2021; Chen et al., 2024), consistent with patterns observed in the core microbiome analysis. Conversely, both amended houses exhibited lower relative abundance of the halophilic genera *Natribacillus* and *Gracilibacillus*, potentially reflecting differences in litter salinity associated with reduced NaHSO₄ application. These associations suggest that variation in litter acidification and ionic conditions may have contributed to the observed microbiome differences, although their respective contributions were not determined. Both amended houses also exhibited lower relative abundance of *Staphylococcus*, a genus commonly present in poultry litter and on bird skin and mucosa (Lowder et al., 2009).

Several limitations should be considered when interpreting the findings of the present study. Each management regimen was represented by a single commercial house, and although six composite samples were collected per house, these captured within-house spatial variation rather than independent treatment replication. Consequently, observed microbiome differences cannot be attributed to litter management independently of house-specific factors. Although houses were selected based on comparable management and productivity histories, the absence of baseline sampling precluded assessment of initial microbiome differences. Sampling was also limited to a single time point following bird removal, preventing evaluation of microbial succession throughout the flock cycle (Johnson et al., 2021; Kubasova et al., 2022). Furthermore, the absence of concurrent litter physicochemical measurements (e.g., pH, moisture, and NH₃) and bird performance data limited interpretation of the functional and production-related implications of the observed compositional differences. Because the postbiotic was administered alongside a reduced NaHSO₄ rate, the individual contributions of these management components also could not be distinguished. Future studies should incorporate independent house replication, baseline and longitudinal sampling, concurrent physicochemical and performance measurements, and factorial designs to evaluate the respective contributions of postbiotic application and acidifier rate.

## Conclusion

This study characterized the litter microbiome of commercial broiler houses under contrasting litter management regimens. Litter from houses receiving a postbiotic amendment with reduced NaHSO₄ differed in community composition from that of a house receiving the standard NaHSO₄ rate, while α diversity did not differ. Both amended houses showed consistent differences in core microbiome membership and in the relative abundance of several genera, including higher *Bifidobacterium* and lower *Staphylococcus*. Although these house-level differences cannot be distinguished from house-specific variation, they identify candidate taxa and community features for evaluation in future trials. Replicated trials incorporating baseline and longitudinal sampling, litter physicochemistry, and flock performance will help establish whether these patterns are reproducible and determine their practical significance.

## Data availability

The FASTQ files associated with this work have been deposited in the NCBI Sequence Read Archive database under BioProject PRJNA1332267.

## Author contributions

Formal analysis, B.H.; interpretation, E.G.O., G.G., S.C.R. and B.H.; visualization, B.H.; writing—original draft preparation, E.G.O. and B.H.; writing—review and editing, E.G.O., G.G., S.C.R. and B.H. All authors have read and agreed to the final version of the manuscript and agree to be accountable for all aspects of the work.

## References

Bolan, N. S., Szogi, A. A., Chuasavathi, T., Seshadri, B., Rothrock Jr, M. J., & Panneerselvam, P. (2010). Uses and management of poultry litter. World’s Poultry Science Journal, 66(4), 673–698. 10.1017/S0043933910000656

Olson, E. G., Hale, B. M., & Ricke, S. C. (2026). The Living Floor: Poultry Litter as an Event-Driven Interface Between Bird Health, Environment, and Microbial Communities. Poultry Science, In Press. 10.1016/j.psj.2026.108083

Ricke, S. C., Olson, E. G., Priddle, C. T., Peckenschneider, M. A., & Hale, B. M. (2026a). Chapter 7. An overview of poultry litter microbial ecology and application of microbiome analyses. In: Sunil Thomas, (Ed.) Animal Microbiomes, CRC Press, Taylor & Francis Group, LLC. Boca Raton, FL. pp 149–156. doi.10.1201/9781003638568-7.

Ritz, C. W., Fairchild, B. D., & Lacy, M. P. (2004). Implications of ammonia production and emissions from commercial poultry facilities: A review. Journal of Applied Poultry Research, 13(4), 684–692. 10.1093/japr/13.4.684

Rothrock, M. J., Cook, K. L., Warren, J. G., Eiteman, M. A., & Sistani, K. (2010). Microbial mineralization of organic nitrogen forms in poultry litters. Journal of Environmental Quality, 39(5), 1848–1857. 10.2134/jeq2010.0024

Johnson, T. M., Fairchild, B., & Ritz, C. W. (2011). The use of sodium bisulfate as a best management practice for reducing ammonia emissions from poultry manures. NPWMS Proceedings.

Toppel, K., Kaufmann, F., Schön, H., Gauly, M., & Andersson, R. (2019). Effect of pH-lowering litter amendment on animal-based welfare indicators and litter quality in a European commercial broiler husbandry. Poultry Science, 98(3), 1181–1189. 10.3382/ps/pey489

de Toledo, T. D. S., Roll, A. A. P., Rutz, F., Dallmann, H. M., Dai Pra, M. A., Leite, F. P. L., & Roll, V. F. B. (2020). An assessment of the impacts of litter treatments on the litter quality and broiler performance: A systematic review and meta-analysis. PLoS One, 15(5), e0232853. 10.1371/journal.pone.0232853

Johnson, J., Zwirzitz, B., Oladeinde, A., Milfort, M., Looft, T., Chai, L., … & Aggrey, S. E. (2021). Succession patterns of the bacterial community in poultry litter after bird removal and sodium bisulfate application Journal of Environmental Quality, 50(4), 923–933. 10.1002/jeq2.20248

Williams, Z. T., Blake, J. P., & Macklin, K. S. (2012). The effect of sodium bisulfate on Salmonella viability in broiler litter. Poultry Science, 91(9), 2083–2088. 10.3382/ps.2011-01976

Joerger, R. D., Ganguly, A., de Los Santos, M., & Li, H. (2020). Effect of sodium bisulfate amendments on bacterial populations in broiler litter. Poultry Science, 99(11), 5560–5571. 10.1016/j.psj.2020.08.013

Callaway, T.R. and S.C. Ricke (Eds.), 2023. Direct Fed Microbials/Prebiotics for Animals: Science and Mechanisms of Action, 2nd Edition. Springer Science, New York, NY, 348 pp.

Olson, E. G., Hale, B. M., Chatman, C. C., Mantovani, H. C., Majumder, E. L. W., & Ricke, S. C. (2025). Gut microbiota and advances in microbiome sequencing-based technologies: opportunities for potential biologics discovery in meat animals. Meat and Muscle Biology, 9(1). 10.22175/mmb.20022

Ricke, S.C., Rogers, A.E., Olson, E.G., & Hale, B.M. (2026b). Bacterial pathogens occurring in poultry litter. Poultry Sci. 105: 108046. 10.1016/j.psj.2026.108046.

Pedroso, A. A., Hurley-Bacon, A. L., Zedek, A. S., Kwan, T. W., Jordan, A. P., Avellaneda, G., … & Lee, M. D. (2013). Can probiotics improve the environmental microbiome and resistome of commercial poultry production?. International Journal of Environmental Research and Public Health, 10(10), 4534–4559. 10.3390/ijerph10104534

Gilchrist, M. J., Greko, C., Wallinga, D. B., Beran, G. W., Riley, D. G., & Thorne, P. S. (2007). The potential role of concentrated animal feeding operations in infectious disease epidemics and antibiotic resistance. Environmental Health Perspectives, 115(2), 313–316. 10.1289/ehp.8837

Olson, E. G., & Hale, B. (2026). Avian Gastrointestinal Microbiota and Its Role in Poultry Health. In Encyclopedia of Livestock Medicine for Large Animal and Poultry Production (pp. 1–6). Cham: Springer Nature Switzerland. 10.1007/978-3-031-52133-1_378-1

Locatelli, A., Hiett, K. L., Caudill, A. C., & Rothrock Jr, M. J. (2017). Do fecal and litter microbiomes vary within the major areas of a commercial poultry house, and does this affect sampling strategies for whole-house microbiomic studies?. Journal of Applied Poultry Research, 26(3), 325–336. 10.3382/japr/pfw076

Caporaso, J. G., Lauber, C. L., Walters, W. A., Berg-Lyons, D., Lozupone, C. A., Turnbaugh, P. J., … & Knight, R. (2011). Global patterns of 16S rRNA diversity at a depth of millions of sequences per sample. Proceedings of the National Academy of Sciences, 108(supplement_1), 4516-4522. 10.1073/pnas.1000080107

Walters, W. A., Caporaso, J. G., Lauber, C. L., Berg-Lyons, D., Fierer, N., & Knight, R. (2011). PrimerProspector: de novo design and taxonomic analysis of barcoded polymerase chain reaction primers. Bioinformatics, 27(8), 1159–1161. 10.1093/bioinformatics/btr087

Martin, M. (2011). Cutadapt removes adapter sequences from high-throughput sequencing reads. *EMBnet*. Journal, 17(1), 10–12. 10.14806/ej.17.1.200

Callahan, B. J., McMurdie, P. J., Rosen, M. J., Han, A. W., Johnson, A. J. A., & Holmes, S. P. (2016). DADA2: High-resolution sample inference from Illumina amplicon data. Nature Methods, 13(7), 581–583. 10.1038/nmeth.3869

Bolyen, E., Rideout, J. R., Dillon, M. R., Bokulich, N. A., Abnet, C. C., Al-Ghalith, G. A., … & Caporaso, J. G. (2019). Reproducible, interactive, scalable and extensible microbiome data science using QIIME 2. Nature Biotechnology, 37(8), 852–857. 10.1038/s41587-019-0209-9

Quast, C., Pruesse, E., Yilmaz, P., Gerken, J., Schweer, T., Yarza, P., & Glöckner, F. O. (2012). The SILVA ribosomal RNA gene database project: improved data processing and web-based tools. Nucleic Acids Research, 41(D1), D590–D596. 10.1093/nar/gks1219

R Core Team. (2024). R: A language and environment for statistical computing. R Foundation for Statistical Computing, Vienna, Austria. https://www.R-project.org/

Posit team. (2024). RStudio: Integrated development environment for R. Posit Software, PBC, Boston, MA. http://www.posit.co/

Bisanz, J. E. (2018). qiime2R: Importing QIIME2 artifacts and associated data into R sessions. Version 0.99, 3.

McMurdie, P. J., & Holmes, S. (2013). phyloseq: an R package for reproducible interactive analysis and graphics of microbiome census data. PloS One, 8(4), e61217. 10.1371/journal.pone.0061217

Mikryukov, V. (2020). metagMisc: Miscellaneous functions for metagenomic analysis.

Good, I. J. (1953). The population frequencies of species and the estimation of population parameters. Biometrika, 40(3-4), 237–264. 10.1093/biomet/40.3-4.237

Good, I. J., & Toulmin, G. H. (1956). The number of new species, and the increase in population coverage, when a sample is increased. Biometrika, 43(1-2), 45–63. 10.1093/biomet/43.1-2.45

Chao, A., & Jost, L. (2012). Coverage-based rarefaction and extrapolation: standardizing samples by completeness rather than size. Ecology, 93(12), 2533–2547. 10.1890/11-1952.1

Dixon, P. (2003). VEGAN, a package of R functions for community ecology. Journal of Vegetation Science, 14(6), 927–930. 10.1111/j.1654-1103.2003.tb02228.x

Wickham, H., & Sievert, C. (2009). ggplot2: elegant graphics for data analysis (Vol. 10, pp. 978–0). New York: springer. 10.18637/jss.v077.b02

Paradis, E., Claude, J., & Strimmer, K. (2004). APE: analyses of phylogenetics and evolution in R language. Bioinformatics, 20(2), 289–290. 10.1093/bioinformatics/btg412

Yu, G., Smith, D. K., Zhu, H., Guan, Y., & Lam, T. T. Y. (2017). ggtree: an R package for visualization and annotation of phylogenetic trees with their covariates and other associated data. Methods in Ecology and Evolution, 8(1), 28–36. 10.1111/2041-210X.12628

Shannon, C. E. (1948). A mathematical theory of communication. The Bell System Technical Journal, 27(3), 379–423. 10.1002/j.1538-7305.1948.tb01338.x

Pielou, E. C. (1966). The measurement of diversity in different types of biological collections. Journal of Theoretical Biology, 13, 131–144. 10.1016/0022-5193(66)90013-0

Lahti, L., & Shetty, S. (2018). Introduction to the microbiome R package. *Preprint at* https://microbiome.github.io/tutorials.

Faith, D. P. (1992). Conservation evaluation and phylogenetic diversity. Biological Conservation, 61(1), 1–10. 10.1016/0006-3207(92)91201-3

Kembel, S. W., Cowan, P. D., Helmus, M. R., Cornwell, W. K., Morlon, H., Ackerly, D. D., … & Webb, C. O. (2010). Picante: R tools for integrating phylogenies and ecology. Bioinformatics, 26(11), 1463–1464. 10.1093/bioinformatics/btq166

Levene, H. (1960). Robust tests for equality of variances. Contributions to Probability and Statistics, 278–292.

Fox, J., Weisberg, S., Adler, D., Bates, D., Baud-Bovy, G., Ellison, S., & Heiberger, R. (2012). Package ‘car’. Vienna: R Foundation for Statistical Computing, 16(332), 333.

Shapiro, S. S., & Wilk, M. B. (1965). An analysis of variance test for normality (complete samples). Biometrika, 52(3-4), 591–611. 10.1093/biomet/52.3-4.591

Paulson, J. N., Stine, O. C., Bravo, H. C., & Pop, M. (2013). Differential abundance analysis for microbial marker-gene surveys. Nature Methods, 10(12), 1200–1202. 10.1038/nmeth.2658

Bray, J. R., & Curtis, J. T. (1957). An ordination of the upland forest communities of southern Wisconsin. Ecological Monographs, 27(4), 326–349. 10.2307/1942268

Jaccard, P. (1908). Nouvelles recherches sur la distribution florale. Bull. Soc. Vaud. Sci. Nat., 44, 223–270.

Lozupone, C., & Knight, R. (2005). UniFrac: a new phylogenetic method for comparing microbial communities. Applied and Environmental Microbiology, 71(12), 8228–8235. 10.1128/AEM.71.12.8228-8235.2005

Martinez Arbizu, P. (2020). pairwiseAdonis: Pairwise multilevel comparison using adonis. R package version 0.4, 1.

Gu, Z., Eils, R., & Schlesner, M. (2016). Complex heatmaps reveal patterns and correlations in multidimensional genomic data. Bioinformatics, 32(18), 2847–2849. 10.1093/bioinformatics/btw313

Mejia, G., Jara-Servin, A., Hernández-Álvarez, C., Romero-Chora, L., Peimbert, M., Cruz-Ortega, R., & Alcaraz, L. D. (2025). Rhizosphere microbiome influence on tomato growth under low-nutrient settings. FEMS Microbiology Ecology, 101(3), fiaf019. 10.1093/femsec/fiaf019

Yan, L., & Yan, M. L. (2021, June). Package “ggvenn.”.

Mallick, H., Rahnavard, A., McIver, L. J., Ma, S., Zhang, Y., Nguyen, L. H., … & Huttenhower, C. (2021). Multivariable association discovery in population-scale meta-omics studies. PLoS Computational Biology, 17(11), e1009442. 10.1371/journal.pcbi.1009442

Benjamini, Y., & Hochberg, Y. (1995). Controlling the false discovery rate: a practical and powerful approach to multiple testing. Journal of the Royal Statistical Society: series B (Methodological), 57(1), 289–300. 10.1111/j.2517-6161.1995.tb02031.x

Do, A. D. T., Lozano, A., Van Laar, T. A., Mero, R., Lopez, C., Hisasaga, C., … & Tarrant, K. J. (2024). Evaluating microbiome patterns, microbial species, and leg health associated with reused litter in a commercial broiler barn. Journal of Applied Poultry Research, 33(4), 100490. 10.1016/j.japr.2024.100490

Neu, A. T., Allen, E. E., & Roy, K. (2021). Defining and quantifying the core microbiome: challenges and prospects. Proceedings of the National Academy of Sciences, 118(51), e2104429118. 10.1073/pnas.2104429118

Jiao, S., Chen, W., & Wei, G. (2022). Core microbiota drive functional stability of soil microbiome in reforestation ecosystems. Global Change Biology, 28(3), 1038–1047. 10.1111/gcb.16024

Shi, C., Guo, L., Yang, H., Leng, X., Deng, P., Bi, J., & Wang, Y. (2023). Effect of dietary supplementation of Ganoderma lucidum residue on growth performance, immune organ index and faecal microbial community diversity in broiler chickens. Research Square. 10.21203/rs.3.rs-2976135/v1

Cressman, M. D., Yu, Z., Nelson, M. C., Moeller, S. J., Lilburn, M. S., & Zerby, H. N. (2010). Interrelations between the microbiotas in the litter and in the intestines of commercial broiler chickens. Applied and Environmental Microbiology, 76(19), 6572–6582. 10.1128/AEM.00180-10

He, Z., Liu, R., Wang, M., Wang, Q., Zheng, J., Ding, J., … & Zhao, G. (2023). Combined effect of microbially derived cecal SCFA and host genetics on feed efficiency in broiler chickens. Microbiome, 11(1), 198. 10.1186/s40168-023-01627-6

Xu, P., Liu, C., Ding, H., Chen, P., Fan, X., Wang, X., & Xiao, Y. (2025). Effect of bio-fermented distillers grain on growth, intestines, and caecal microbial community in broilers. Fermentation, 11(3), 118. 10.3390/fermentation11030118

Grant, A. Q., Gay, C. G., & Lillehoj, H. S. (2018). Bacillus spp. as direct-fed microbial antibiotic alternatives to enhance growth, immunity, and gut health in poultry. Avian Pathology, 47(4), 339–351. 10.1080/03079457.2018.1464117

Wu, Y., Wang, B., Zeng, Z., Liu, R., Tang, L., Gong, L., & Li, W. (2019). Effects of probiotics Lactobacillus plantarum 16 and Paenibacillus polymyxa 10 on intestinal barrier function, antioxidative capacity, apoptosis, immune response, and biochemical parameters in broilers. Poultry Science, 98(10), 5028–5039. 10.3382/ps/pez226

Halder, N., Sunder, J., De, A. K., Bhattacharya, D., & Joardar, S. N. (2024). Probiotics in poultry: a comprehensive review. The Journal of Basic and Applied Zoology, 85(1), 23. 10.1186/s41936-024-00379-5

Kairmi, S. H., Abdelaziz, K., Spahany, H., Astill, J., Trott, D., Wang, B., & Sharif, S. (2024). Intestinal microbiome profiles in broiler chickens raised without antibiotics exhibit altered microbiome dynamics relative to conventionally raised chickens. Plos One, 19(4), e0301110. 10.1371/journal.pone.0301110

Kubasova, T., Faldynova, M., Crhanova, M., Karasova, D., Zeman, M., Babak, V., & Rychlik, I. (2022). Succession, replacement, and modification of chicken litter microbiota. Applied and Environmental Microbiology, 88(24), e01809–22. 10.1128/aem.01809-22

Wang, H., Shankar, V., & Jiang, X. (2022). Compositional and functional changes in microbial communities of composts due to the composting-related factors and the presence of Listeria monocytogenes. Microbiology Spectrum, 10(4), e01845–21. 10.1128/spectrum.01845-21

Valečková, E., Sun, L., Wang, H., Dube, F., Ivarsson, E., Kasmaei, K. M., … & Wall, H. (2023). Intestinal colonization with Campylobacter jejuni affects broiler gut microbiota composition but is not inhibited by daily intake of Lactiplantibacillus plantarum. Frontiers in Microbiology, 14, 1205797. 10.3389/fmicb.2023.1205797

Dumas, M. D., Polson, S. W., Ritter, D., Ravel, J., Gelb Jr, J., Morgan, R., & Wommack, K. E. (2011). Impacts of poultry house environment on poultry litter bacterial community composition. PLoS One, 6(9), e24785. 10.1371/journal.pone.0024785

Poehlein, A., Yutin, N., Daniel, R., & Galperin, M. Y. (2017). Proposal for the reclassification of obligately purine-fermenting bacteria *Clostridium acidurici* (Barker 1938) and *Clostridium purinilyticum* (Dürre et al. 1981) as *Gottschalkia acidurici* gen. nov. comb. nov. and *Gottschalkia purinilytica* comb. nov. and of *Eubacterium angustum* (Beuscher and Andreesen 1985) as *Andreesenia angusta* gen. nov. comb. nov. in the family Gottschalkiaceae fam. nov. International Journal of Systematic and Evolutionary Microbiology, 67(8), 2711–2719. 10.1099/ijsem.0.002008

Lu, J., Sanchez, S., Hofacre, C., Maurer, J. J., Harmon, B. G., & Lee, M. D. (2003). Evaluation of broiler litter with reference to the microbial composition as assessed by using 16S rRNA and functional gene markers. Applied and Environmental Microbiology, 69(2), 901–908. 10.1128/AEM.69.2.901-908.2003

Chen, L., Han, M., Xu, J., Cao, Z., Chen, W., Jing, B., … & Wen, X. (2024). Firmicutes primarily drive odor emission profiles in poultry manure treatments. Poultry Science, 103(11), 104250. 10.1016/j.psj.2024.104250

Lowder, B. V., Guinane, C. M., Ben Zakour, N. L., Weinert, L. A., Conway-Morris, A., Cartwright, R. A., … & Fitzgerald, J. R. (2009). Recent human-to-poultry host jump, adaptation, and pandemic spread of Staphylococcus aureus. Proceedings of the National Academy of Sciences, 106(46), 19545–19550. 10.1073/pnas.0909285106

